# Genomic and geographic footprints of differential introgression between two highly divergent fish species

**DOI:** 10.1101/225201

**Authors:** Ahmed Souissi, François Bonhomme, Manuel Manchado, Lilia Bahri-Sfar, Pierre-Alexandre Gagnaire

## Abstract

Investigating variation in gene flow across the genome between closely related species is important to understand how reproductive isolation builds up during the speciation process. An efficient way to characterize differential gene flow is to study how the genetic interactions that take place in hybrid zones selectively filter gene exchange between species, leading to heterogeneous genome divergence. In the present study, genome-wide divergence and introgression patterns were investigated between two sole species, *Solea senegalensis* and *Solea aegyptiaca*, using a restriction-associated DNA sequencing (RAD-Seq) approach to analyze samples taken from a transect spanning the hybrid zone. An integrative approach combining geographic and genomic clines methods with an analysis of individual locus introgression taking into account the demographic history of divergence inferred from the joint allele frequency spectrum was conducted. Our results showed that only a minor fraction of the genome can still substantially introgress between the two species due to genome-wide congealing. We found multiple evidence for a preferential direction of introgression in the *S. aegyptiaca* genetic background, indicating a possible recent or ongoing movement of the hybrid zone. Deviant introgression signals found in the opposite direction supported that the Mediterranean populations of *S. senegalensis* could have benefited from adaptive introgression. Our study thus illustrates the varied outcomes of genetic interactions between divergent gene pools that recently met after a long history of divergence.

## INTRODUCTION

Reproductive isolation and hybridisation are antinomic but related processes that are tuned by the same evolutionary forces. Natural hybridisation between two divergent lineages occurs as long as reproductive isolation is not totally established (Dobzhansky, 1937; Mayr, 1942; Coyne and Orr, 2004). The resulting exchanges of genetic material beyond the first hybrid generation occur through the production of backcross and recombinant genotypes that are exposed to selection, allowing neutral and advantageous alleles to spread among divergent lineages but preventing gene flow at the loci involved in reproductive isolation (Barton, 1979). Hybrid zones, the geographic areas where hybridisation occurs, therefore act as selective filters which allow us to unravel the strength of reproductive barriers and their impact on the patterns and dynamics of gene exchanges at various stage of speciation (Barton and Hewitt, 1985; Barton and Bengtsson, 1986; Hewitt, 1988; Martinsen *et al*., 2001).

The semipermeable nature of species boundaries was first evidenced from differential introgression patterns in hybrid zones (Harrison, 1990), that is, variation among loci in the level of incorporation of alleles from one lineage into the other (Payseur, 2010; Harrison and Larson, 2016). Introgression at a given neutral marker depends on the antagonistic effects between counter-selection on a nearby selected locus and recombination with that barrier locus (Barton and Bengtsson, 1986). Therefore, differential gene flow across the genome reflects both recombinational distance to a barrier locus and selection acting on that locus. Different methods have been developed to detect individual loci with introgression behaviours departing from the genome-wide average. This includes the analysis of spatial allele frequency patterns in a transect spanning the hybrid zone, or variation in ancestry proportion at individual loci compared to genome-wide expectations in admixed genotypes (see Payseur 2010 for a review).

The geographic cline method is a powerful approach to analyse the relationship between allele frequency and geographic distance to the center of a hybrid zone (Endler, 1977; Barton and Hewitt, 1985; Barton and Gale, 1993; Porter *et al*., 1997; Teeter *et al*., 2008). Fitting a cline model to observed allele frequency patterns allows inferring fundamental parameters such as cline center, dispersal and the strength of selection. Therefore, important information concerning the balance between migration and selection can be obtained using the geographic cline method, such as the identification of which loci tend to introgress neutrally and which do not. The genomic cline method is a related approach developed to deal with hybrid zones such as mosaic hybrid zones that, unlike clinal hybrid zones, are not spatially structured, (Harrison and Rand, 1989; Bierne *et al*., 2003; Larson *et al*., 2013). The change in allele frequency at individual loci is analysed along a gradient of genomic admixture instead of a spatial gradient, enabling a comparison of individual locus introgression patterns with regards to the genome-wide average pattern of admixture (Gompert and Buerkle, 2009, 2011; Fitzpatrick, 2013). As for geographic clines, this genomic cline method also provides a quantitative assessment of the excess of ancestry to a given parental species, as well as the rate of change in allele frequency from one species to the other along the admixture gradient. Although both approaches are useful to document the effect of selection and recombination on differential introgression in admixed genotypes, they do not specifically account for the demographic history of the parental species (Payseur and Rieseberg, 2016).

An alternative approach that partly overcomes this limitation is the use of demo-genetic models that account for varying rates of introgression among loci during the divergence history. Such methods for reconstructing the history of gene flow between semi-isolated lineages have been developed within Bayesian (Sousa *et al*., 2013), approximate Bayesian computation (Roux *et al*., 2013, 2016), or approximate likelihood frameworks (Tine *et al*., 2014; Le Moan *et al*., 2016; Rougeux *et al*., 2017). Using summary statistics like the joint allelic frequency spectrum, which depicts correlations in allele frequencies between lineages outside the hybrid zone, they capture variable behaviours among loci and allow quantifying the degree of semi-permeability reflecting the overall balance between gene flow and selection. These methods have the advantage of specifically accounting for the history of gene flow during divergence, using contrasted speciation scenarios such as primary differentiation, ancient migration or secondary contact. However, they do not provide information about individual locus behaviour as the cline methods do. Here, we pushed these inference methods one step further in order to assess the probability for a given locus to belong to one of two categories: (i) loci with a reduced effective migration rate due to selection and linkage, and (ii) loci which can readily introgress.

The flatfish species *Solea senegalensis* and *Solea aegyptiaca* are two economically important species in the Mediterranean basin that hybridise along the northern Tunisian coasts, where they form a hybrid zone (She *et al*., 1987). Mitochondrial divergence between them is high (~ 2%) if compared to other vertebrate species-pairs that also hybridise in nature, such as the house mice *Mus m. musculus* and *M. m. domesticus* (0.3%, based on sequences retrieved from Genebank) or Atlantic and Mediterranean lineages of *Dicentrarchus labrax* (0.7%, Tine *et al*., 2014). Based on an analysis of spatial allele frequency patterns at a dozen of allozymes and intronic loci, Ouanes *et al*., (2011) proposed that the hybrid zone between *S. senegalensis* and *S. aegyptiaca* was centred in Bizerte lagoon acting as a non-stable unimodal tension zone stemming from a secondary contact. They also suggested that the zone could have undergone a recent expansion. Recently, Souissi *et al*., (2017) showed the existence of morphological transgressions within the contact zone, possibly indicating a reduced fitness of recombined compared to parental genotypes. However, the low number of markers in these latter studies could not provide a clear description of the genetic exchanges across the genome of these two species, nor quantifying the strength of the forces maintaining the incomplete reproductive isolation between them.

In the present work, high-throughput genotyping using RAD sequencing (Baird *et al*., 2008) was carried out in individuals sampled at both sides of the hybrid zone, with a particular effort in the contact zone itself. By combining three different methods (genomic and geographic cline analysis and historical demographic inference) exploiting different aspects of the data, we provide new insights on the history of divergence between *S. senegalensis* and *S. aegyptiaca*, and a genome-wide description of varied patterns of introgression attributed to recent or ongoing movement of the hybrid zone.

## MATERIALS AND METHODS

### Sampling

A total of 153 samples were collected from 10 locations spanning the distribution range of *Solea senegalensis* and *S. aegyptiaca* from Senegal to Egypt (Fig.1). This sampling strategy aimed at covering the geographical distribution of both species, including a detailed transect of their natural hybrid zone in Tunisia. Five sampling locations were collected throughout the *S. senegalensis* parental zone, two in the Atlantic Ocean (10 individuals from Dakar in Senegal and 16 from the Gulf of Cadiz in Spain) and three in the Western Mediterranean Sea (15 individuals from Annaba and 8 from Mellah Lagoon in Algeria, and 10 from Tabarka in Tunisia). Three locations were sampled across the geographical range of *S. aegyptiaca* in the Eastern Mediterranean Sea (13 individuals from Kerkennah Island and 15 from El Biban lagoon in Tunisia, and 20 samples from Bardawill lagoon in Egypt). Sampling size was locally increased in the Tunisian region where both species coexist and hybridisation has been reported (Bizerte lagoon *n* = 29 and Gulf of Tunis *n* = 25). Finally, a total of 7 individuals belonging to the closely related species *S. solea* were sampled in Tunisia to provide an outgroup species for the orientation of ancestral and derived alleles in *S. senegalensis* and *S. aegyptiaca*.

**Figure 1:**
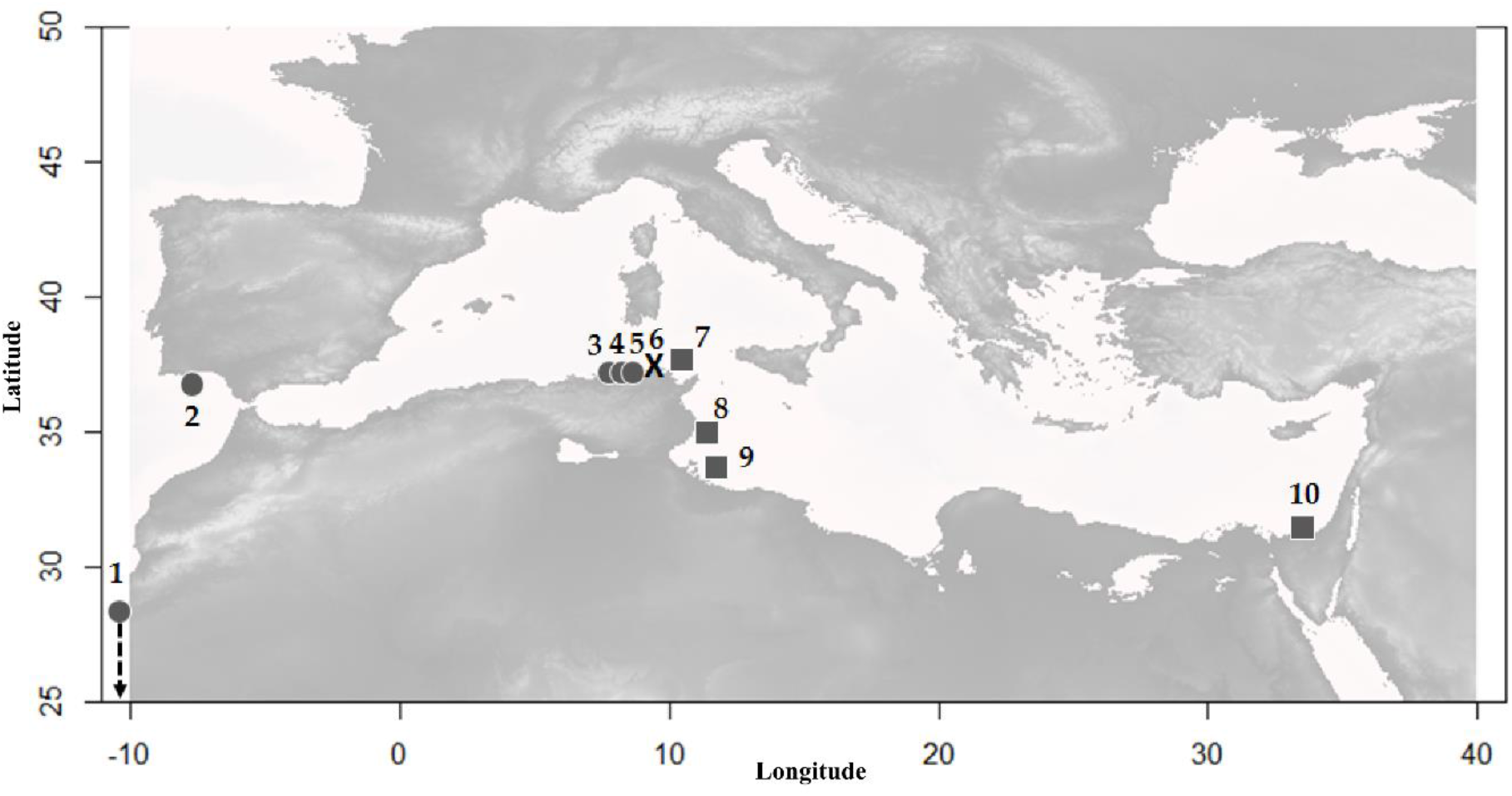
Map of sampling locations: **1)** Dakar, **2)** Cadiz, **3)** Annaba, **4)** Mellah Lagoon, **5)** Tabarka, **6)** Bizerte Lagoon, **7)** Gulf of Tunis, **8)** Kerkennah Islands, **9)** El Bibane Lagoon, **10)** Bardawill Lagoon.

### RAD library preparation and sequencing

Whole genomic DNA was extracted from fin clips using the DNeasy Blood & Tissue kit (Qiagen). The presence of high molecular weight DNA was checked on a 1% agarose gel, and double-stranded DNA concentration was quantified using Qubit 2.0 and standardized to 25 ng per μl. Restriction-Associated DNA (RAD) library construction followed a modified version of the original single-end RAD-seq protocol (Baird *et al*., 2008). Briefly, 1 μg of genomic DNA from each individual was digested using the restriction enzyme *Sbf*I-HF (NEB), and ligated to one of 32 unique molecular barcodes of 5 to 6 bp. Ligated products were then combined in equimolar proportions into five RAD libraries, each made of a multiplex of 32 individuals originating from various localities. Each library was finally sequenced in 101 bp single read mode on a separate lane of an Illumina HiSeq2500 sequencer, at the sequencing platform ‘Génomique Intégrative et Modélisation des Maladies Métaboliques’ (UMR 8199, Lille, France).

### Bioinformatic analyses

Raw reads were de-multiplexed based on individual barcode information, end-clipped to 95 pb and quality filtered using the program *process_radtags* from the *stacks* pipeline with default settings (Catchen *et al*., 2013). Retained read were then aligned to a draft assembly of the *S. senegalensis* genome (98,590 scaffolds, total length 740 Mb, N50 contig length 10,767 bp, Manchado *et al*., 2016) using bowtie 2 v.2.1.0 (Langmead and Salzberg, 2012) with the -- very-sensitive option. In order to take into account both the level of divergence among species and the possibility of introgression, we used a subset of individuals to empirically determine the optimal maximum number of mismatches allowed between aligned reads and the reference genome (m = 7). We then used *pstacks* to call variable positions under the bounded SNP model, setting the upper sequencing error rate to 2.5% and a minimum sequencing depth to 5X per stack. Homologous loci across samples were merged based on their genomic position within scaffolds using *cstacks* to construct a catalogue of loci. Individual stacks were then matched against the catalogue of loci with *sstacks* to determine genotypes. The module *rxstacks* was ran with the --prune_haplo option to exclude poor quality loci using a log-likelihood threshold of −300 (determined empirically). Individual genotypes were finally exported in VCF format using the module *populations*.

We then applied population-specific filters with *VCFtools* (Danecek *et al*., 2011) to remove SNPs showing significant deviation to Hardy–Weinberg equilibrium within *S. senegalensis* and *S. aegyptiaca* samples located outside the hybrid zone, using a *P*-value threshold of 0.01. Next, we excluded loci displaying more than 20% of missing genotypes. Over the 116,385 remaining SNPs, we randomly selected one single SNP for each pair of RAD loci associated with the same restriction site, in order to limit the impact of linkage disequilibrium. Finally, we only retained loci with available sequence data in the outgroup species *S. solea*, which had to be fixed for the polymorphic site found in *S. senegalensis* and *S. aegyptiaca*. This resulted in a final dataset containing 10,758 SNPs.

### Genetic structure and hybridisation

Genetic variation within and among *S. senegalensis* and *S. aegyptiaca* was characterized with a principal component analysis (PCA) performed with the R package *Adegenet* (Jombart, 2008; Jombart and Ahmed, 2011). We then estimated individual admixture proportions from two differentiated genetic clusters using *fastStructure* (Raj *et al*., 2014) with 10^8^ iterations, in order to distinguish parental and admixed genotypes.

### Demo-genetic inference of the divergence history

We used a modified version of the *δaδi* program (Gutenkunst *et al*., 2009) to infer the divergence history between *S. senegalensis* and *S. aegyptiaca*. This method uses the joint allele frequency spectrum (JAFS) between two populations as summary statistics to characterize divergence. *δaδi* fits demographic divergence models to the observed data using a diffusion approximation of the JAFS, enabling to compare different alternative divergence models in a composite likelihood framework. We used seven divergence models developed in a previous study (Tine *et al*., 2014) to determine whether and how gene flow has shaped genome divergence between *S. senegalensis* and *S. aegyptiaca*. The simplest model of strict isolation *(SI)* corresponds to an allopatric divergence scenario in which the ancestral population of effective size *N_A_* splits into two derived populations of size *N_1_* (*S. senegalensis*) and *N_2_* (*S. aegyptiaca*) that evolve without exchanging genes for *T_s_* generations. We then considered three models of divergence including gene flow, either during the entire divergence period (isolation with migration, *IM*), the beginning of divergence (ancient migration, *AM*), or the most recent part of divergence (secondary contact, SC). In these models, migration occurs with potentially asymmetric rates (*m*_12_ and *m*_21_) that are shared across all loci in the genome. We also considered simple extensions of the *IM, AM* and *SC* models that capture the effect of selection by accounting for heterogeneous migration rates among loci (*IM2m, AM2m* and *SC2m*). These semi-permeability models consider that two categories of loci, experiencing different effective migration rates, occur in proportions *P* and 1−*P* in the genome.

The JAFS was obtained by pooling the least introgressed populations from *S. senegalensis* (Dakar, Cadix, Annaba, and Mellah) for species 1 and *S. aegyptiaca* (Kerkennah, L. Bibane, and L. Bardawill) for species 2 (Fig. 2). We used *S. solea* samples as an outgroup species to determine the most parsimonious ancestral allelic state for each SNP in order to generate an unfolded JAFS (Fig. 3a). The size of the JAFS was projected down to 40 sampled chromosomes per species to account for missing data. For each model, we estimated the parameter values that maximize likelihood using two successive simulated annealing (SA) procedures before quasi-Newton (BFGS) optimization (Tine *et al*., 2014). Comparisons among models was made using the Akaike information criterion (AIC) to account for variation in the number of parameters among models. A total of 20 independent runs were used for the optimization of each model. Parameters uncertainties were estimated from non-parametric bootstrapped data using the Godambe information matrix as implemented in *δaδi*. We used 1000 bootstrapped datasets to estimate confidence intervals as the maximum likelihood parameter value ± 1.96×SE.

**Figure 2:**
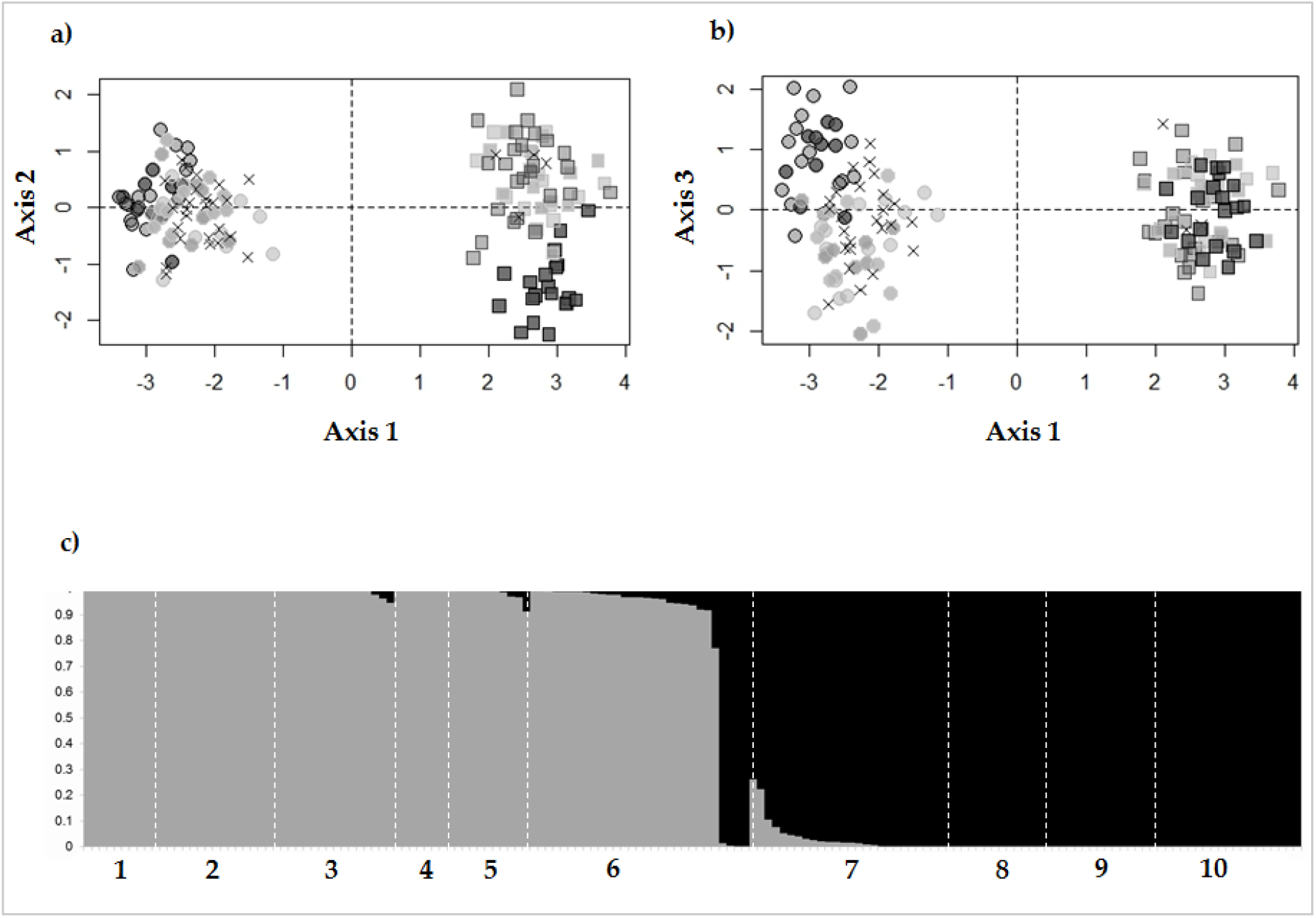
Population genetic structure of *Solea senegalensis* and *Solea aegyptiaca* analysed using 10,758 SNPs. Principal Component Analysis (PCA) of the 153 individuals with panel **a)** representing individual coordinates along PC1 and PC2 axes and panel **b)** showing PC1 and PC3 axes. **c)** *fastStructure* analysis performed with *K*=2 genetic clusters. Numbers indicate the 10 different sampling locations.

### Inferring the probability of locus introgression under the best divergence model

Because the JAFS-based inference of the divergence history does not provide an assessment of introgression probability for each locus separately, we developed an approach to estimate the relative probability of each individual locus to be assigned to one of these two categories: (*i*) loci which can readily introgress between species, and (*ii*) loci experiencing a highly reduced introgression rate due to selection against foreign alleles and linkage. This probability was estimated using the best-fit model identified in the previous section, which was a secondary contact model with variable introgression rates among loci (SC*2m*) (Fig. 3b). The SC*2m* model can be decomposed as a linear combination of two simple models describing gene flow in two different compartments of the genome. The best-fit SC*2m* model estimated that only 5% of the loci can still introgress between species with effective migration rates *m*_1–2_ and *m*_2–1_, whereas the remaining 95% of loci experience a highly reduced introgression rate with effective migration rates *m*’_1–2_ and *m*’_2–1_. We used estimated model parameters to perform coalescent simulations with MSMS (Ewing and Hermisson, 2010) under the secondary contact model (assuming theta=747.202; *N*_1_ =0,818; *N*_2_=1,136; *T*_S_=4,803; *T*_SC_=0,081) using two different conditions for gene flow (assuming either *m*_1–2_=4,607 and *m*_2–1_=0,381, or *m*’_1–2_=0,057 and *m*’_2–1_=0,153) to generate a *free introgression* and a *reduced introgression* rate dataset. One thousand JAFS with 40 sampled chromosomes per species were produced under each model. We then averaged the number of derived alleles within entries across replicates to obtain a single JAFS for both the *free introgression* (ϒ) (Fig. 3c) and the *reduced introgression* (Φ) (Fig. 3d) model. Finally, we used the average number of SNPs per entry (*i,j*) in each JAFS to estimate the probability *P_i,j_* that a SNP with a derived allele count *i* in species 1 and *j* in species 2 can be obtained under the *free introgression* model:

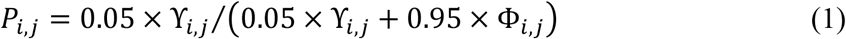

**Figure 3:**
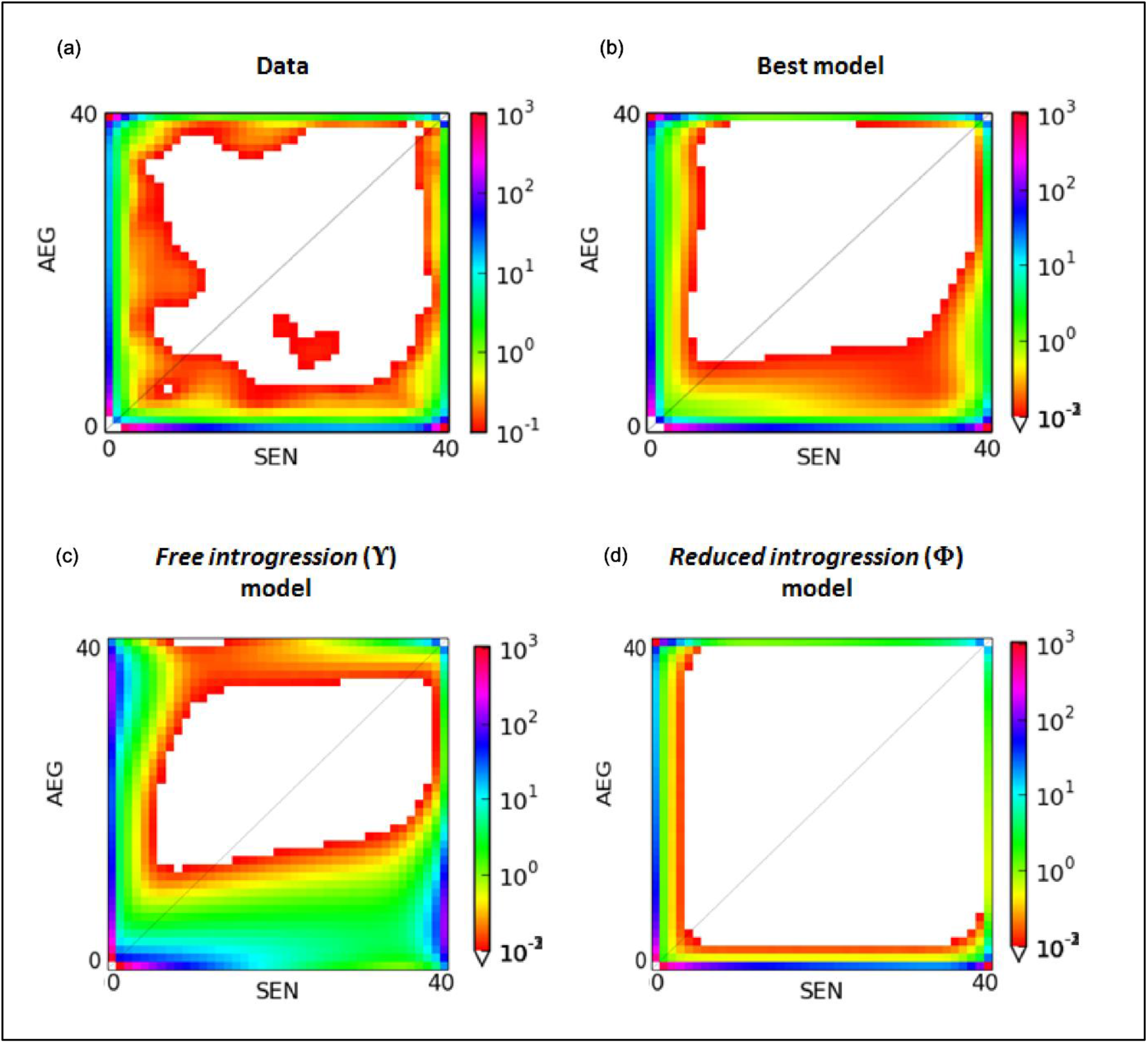
Demographic divergence history of *Solea senegalensis* and *Solea aegyptiaca*. **a)** The observed joint allele frequency spectrum (JAFS) for *S. aegyptiaca* (AEG, y axis) and *S. senegalensis* (SEN, x axis) showing the number of SNPs (coloured scale on the right side of each spectrum) per bin of derived allele counts using 20 individuals per species. **b)** The maximum-likelihood JAFS obtained under the best-fit model of secondary contact with heterogeneous introgression rates (SC*2m* model). **c)** The maximum-likelihood JAFS obtained from *msms* simulations under the free introgression model (ϒ). **d)** The maximum-likelihood JAFS obtained from *msms* simulations under the reduced introgression model Φ).

Where ϒ_*i,j*_ and Φ_*i,j*_ are the average number of SNP predicted in the JAFS entry (*i,j*) under the *free introgression* and *reduced introgression* model, respectively. Every SNP from the real dataset was finally associated to the introgression probability *P_i,j_* given by equation (1) based on its corresponding entry in the JAFS.

### Genomic Clines

The Bayesian Genomic Cline program BGC (Gompert and Buerkle, 2012) was used to quantify individual locus introgression relative to genome-wide introgression. BGC describes the probability of locus-specific ancestry from one parental species given the genome-wide hybrid index. The BGC model considers two principal parameters, called *α* and *β*, which describe locus-specific introgression based on ancestry. Parameter *α* quantifies the change in probability of ancestry relative to a null expectation based on genome-wide hybrid index, that is, the direction of introgression. A positive value of *α* reflects an increase in the probability of ancestry from species one (introgression into *S. aegyptiaca)*, whereas a negative value of *α* reflects an increase in the probability of ancestry from species two (introgression into *S. senegalensis*). Parameter *β* describes the rate of transition in the probability of ancestry from parental population one to parental population two as a function of hybrid index, that is, the amount of introgression. Positives *β* values thus denote a restricted amount of introgression whereas negative *β* values indicate a greater introgression rate compared to the genome-wide average.

We designated outlier loci with respect to parameters *α* and *β* based on the assumption that the genome-wide distributions of locus-specific cline parameters are both centred on zero. Therefore, a locus was considered as an *α* outlier if its posterior estimate of *α* was not contained in the interval bounded by the 0.025 and 0.975 quantiles of 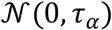 (likewise for β) (Gompert and Buerkle, 2011; Gompert *et al*., 2013).

### Geographic Clines

The geographic cline analysis was carried out in order to link allele frequencies at individual loci with geographic position along a transect spanning the hybrid zone. We used the R package HZAR (Derryberry *et al*., 2014) that fits allele frequency data to classic equilibrium models of geographic clines (Szymura and Barton, 1986) using the MCMC algorithm. We used a model that fits cline center, width, and independent introgression tails using estimated values for minimum (*p*_min_) and maximum (*p*_max_) allele frequencies.

## RESULTS

A total of 833.3 million raw reads were obtained, 89.1% of which were retained after demultiplexing and quality filtering for reference mapping against the *S. senegalensis* draft genome (average number of reads per individual: 3,906,809±1,878,685). Individual genotype calling in Stacks produced a raw VCF file containing 174,490 SNP, from which we retained 10,758 unlinked and oriented SNP that were used for downstream analyses.

### Genetic structure and hybridisation

The Principal Component Analysis (PCA) clearly separated *S. senegalensis* from *S. aegyptiaca* samples along the first PC axis (PC1), which explained 74.2% of the total genotypic variance (Fig. 2a). *S. senegalensis* samples were organised along that axis according to their geographical proximity from the contact zone, following a gradient of genetic similarity to *S. aegyptiaca* increasing from Dakar to Bizerte Lagoon. The two species were found to coexist only in Bizerte Lagoon, with four *S. aegyptiaca* individuals being found among a majority of *S. senegalensis* genotypes. The second principal component (PC2) revealed a weak signal of differentiation (0.73% of associated variance) separating Bardawil lagoon from other samples within *S. aegyptiaca*. The third principal component (PC3, 0.66% of genotypic variance) mostly differentiated Atlantic samples from Dakar and Cadix from the Mediterranean *S. senegalensis* samples (Fig. 2b). The FastStructure analysis (Fig. 2c) confirmed the separation of the two species into different genetic clusters and their coexistence in Bizerte lagoon, as detected from the PCA. The finding of intermediate admixture proportions in some individuals revealed signs of introgressive hybridization between *S. senegalensis* and *S. aegyptiaca* around the contact zone.

### Demo-genetic history of divergence

The *δaδi* analysis showed that the secondary contact model with varying introgression rates along the genome best explained the observed JAFS (SC2*m*, delta AIC > 25). In comparison, the six other models had a significantly lower performance for different reasons. While the strict isolation model (SI) could explain SNPs occupying the outer frame of the JAFS, it could not predict at the same time the presence of loci in the more inner part of the spectrum. The IM, AM and SC models assuming genome-wide homogeneous migration rates better predicted loci occupying the central part of the JASF. However, they underestimated the density of private and highly differentiated SNPs. Finally, the three models including heterogeneous migration rates along the genome (IM2*m*, AM2*m* and SC2*m*) yielded significantly improved fits. However, only the SC2*m* model provided a good prediction for both a high density of highly differentiated SNPs between the two species and a lack of SNPs towards the central part of the spectrum.

Using the best-fit obtained for the SC2*m* model over 20 independent runs to get estimates of model parameters and confidence intervals, we found that the duration of the isolation period between *S. senegalensis* and *S. aegyptiaca* was about 60 times longer than the duration of the secondary contact. A large proportion of the genome (~95%) was associated with relatively small effective migration rates and limited gene flow corresponding to less than one effective migrant per generation in both directions (*N*_1_*m*’_12_ = 0.047, *N*_2_*m*’_21_ = 0.175). By contrast, the effective migration rate was more elevated in the remaining small fraction of the genome (~5%), especially in the direction from *S. senegalensis* to *S. aegyptiaca* where introgression was found 80 times higher than elsewhere in the genome. Introgression was found 12 times lower in direction of *S. senegalensis*, although still more than twice higher than in the remainder (95%) of the genome (Table 1).

**Table 1:**
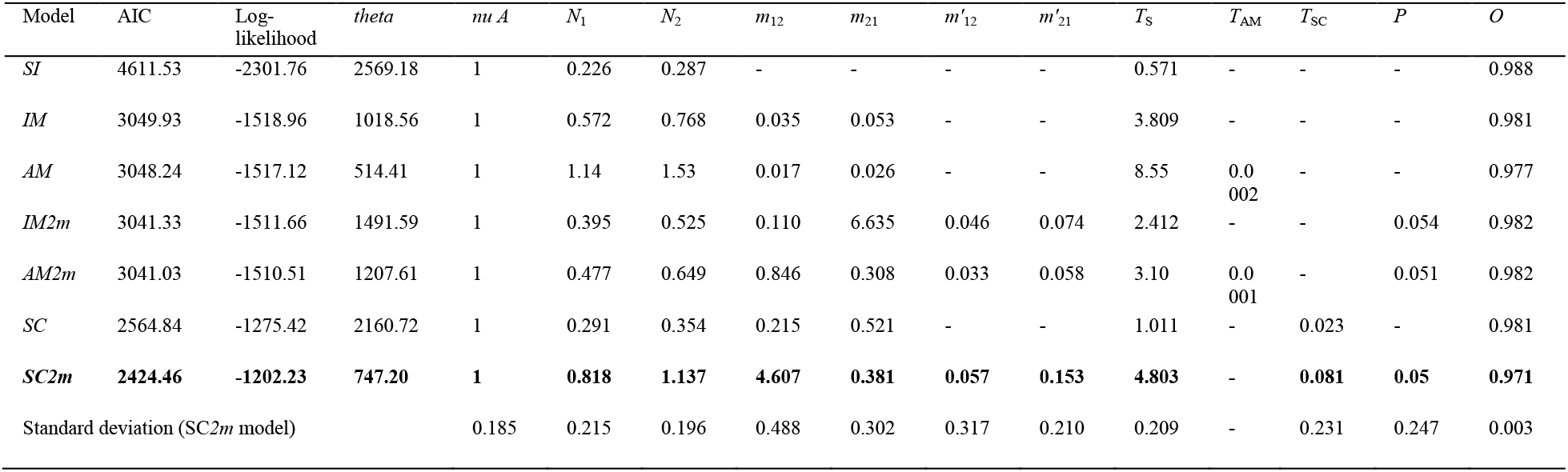
Results of model fitting for seven alternative models of divergence between *S. senegalensis* (SEN) and *S. aegyptiaca* (AEG). In the order of appearance in the table: the model fitted, the Akaike information criterion (AIC), the maximum-likelihood estimate over 20 independent runs (after 3 successive optimization steps: simulated annealing hot, cold and BFGS) and theta parameter for the ancestral population before split. Following are the inferred values for the model parameters (scaled by theta): the effective size of *S. aegyptiaca* (*N*_1_) and *S. senegalensis* (*N*_2_) populations, the effective migration rates for the two categories of loci in the genome (category 1: *m*_12_ and *m*_21_, category 2: *m*’_12_ and *m*’_21_), the duration of the split (*T*_S_) of ancestral migration (*T*_AM_) and secondary contact (*T*_SC_) episodes, and the proportion (*P*) of the genome falling within category 1 (experiencing migration rates *m*_12_ and *m*_21_). The *m*_12_ and *m*’_12_ parameters specify migration rate from population 2 (*S. senegalensis*) into 1 (*S. aegyptiaca*) and reciprocally for *m*_21_ and *m*’_21_. The proportion of correctly orientated SNPs (O) when inferring ancestral allelic states using *S. solea* as an outgroup. *SI*: Strict isolation, *IM*: Isolation with migration, *AM:* Ancient migration, *IM2m*: Isolation with two rates of migration, *AM2m*: Ancient migration with two rates of migration, SC: Secondary contact, SC2m: Secondary contact with two rates of migration.

### Genomic Clines

The genomic cline analysis performed with the BGC program revealed different behaviours among the 10,756 SNPs with respect to the cline parameters *α* and *β*. Considering the locus-wise ancestry shift parameter *α*, we observed that 48% of the loci have an excess of *S. senegalensis* ancestry (negative *α*) whereas only 29% of them have an excess of *S. aegyptiaca* ancestry (positive *α*). The quantile method for outlier detection confirmed that a higher proportion of loci was characterized by an excess of *S. senegalensis* ancestry (43% of negative *α* outliers) compared to *S. aegyptiaca* ancestry (17% of positive *α* outliers). By contrast, the locus-wise slope parameter *β* depicting the rate of transition between species was symmetrically distributed between negative and positive values, with equal proportions of loci showing a decreased (48% of positive *β*) and increased (51% of negative *β*) introgression rates. The proportion of loci showing significant deviation compared to the genomic average amounted to 52%, and were equally distributed among negative (26%) and positive (26%) *β* outliers.

### Geographic clines

The mean position of cline center (C) calculated over all fitted clines was located 12 km to the east of Bizerte lagoon (Fig. 4). Most individual locus clines tended to co-localize at this position, especially for the steepest clines whose centres localised essentially 10 Km or less around the center of the hybrid zone. In general, loci whose individual centres did not coincide with the central part of the hybrid zone harboured less steep clines (Fig. 4)

**Figure 4:**
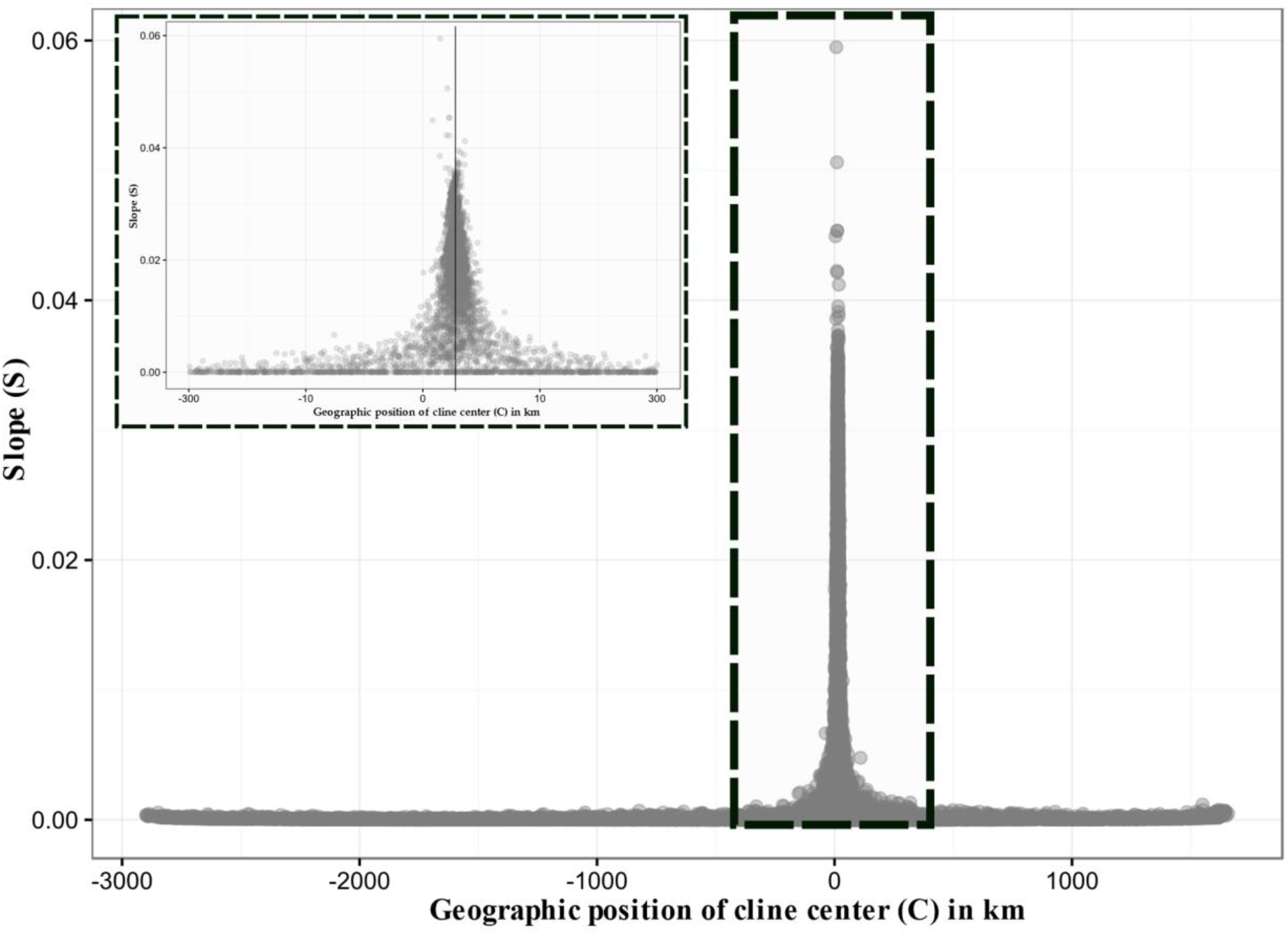
Distribution of geographic cline slope (*S*) for the 10,758 loci as a function of their cline center (*C*). The box represents a zoom in the central part of the contact zone.

### Links between different approaches

Correlation tests were used to investigate whether different approaches that use different aspects of the data tend to produce similar results. First, we evaluated the extent to which an excess of ancestry from a given species relates to a spatial shift of cline center into the other species range. The joint distribution of the genomic cline parameter *α* and geographic cline center *c* (Fig. 5), showed a significant positive correlation on both sides of the hybrid zone. The correlation was however much stronger in the *S. aegyptiaca* side (R^2^_c-α_ = 0.274, *P* < 10^−10^) compared to the *S. senegalensis* side (R^2^_c-α_ = 0.023, *P* < 10^−10^).

We then tested whether the loci exhibiting the most abrupt transition in ancestry between the two species (i.e. a decreased introgression rate) also tended to display the steepest geographic clines. To validate this prediction, we focused on the 26% of positive *β* outliers detected with BGC, and compared *β* values with the slope parameter *S* of the geographic clines. We found a significantly positive correlation (R^2^_S-β_ = 0.27, *P* < 10^−10^) between the two parameters, confirming that loci with a low introgression rate also tend to display steeper geographic clines (Fig. 6).

**Figure 5:**
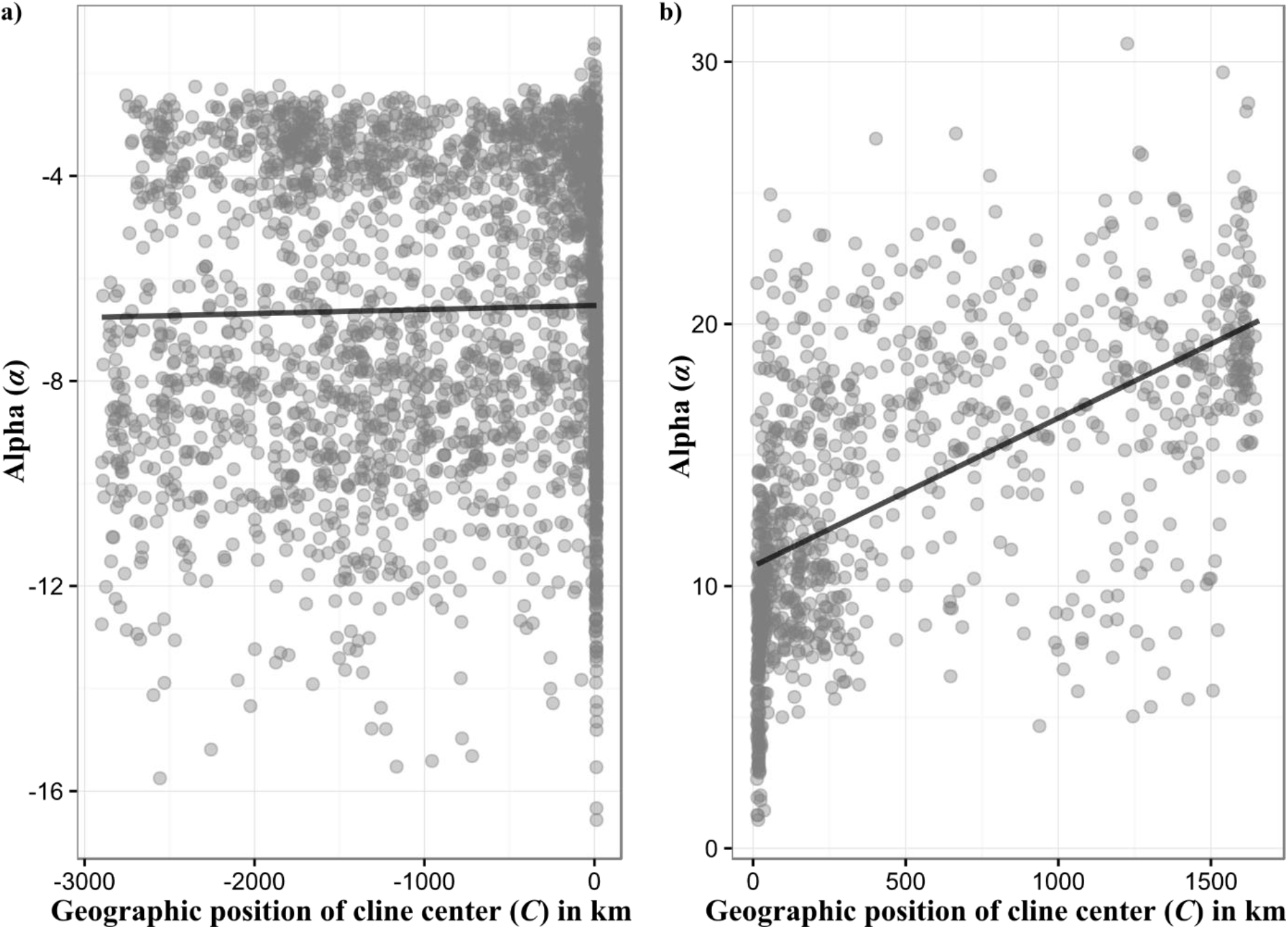
Correlation between genomic cline parameter (*α*) and geographic position of the cline center (*C*) on both sides of the hybrid zone: **a)** in the *Solea senegalensis* geographic range and **b)** in the *Solea aegyptiaca* geographic range.

**Figure 6:**
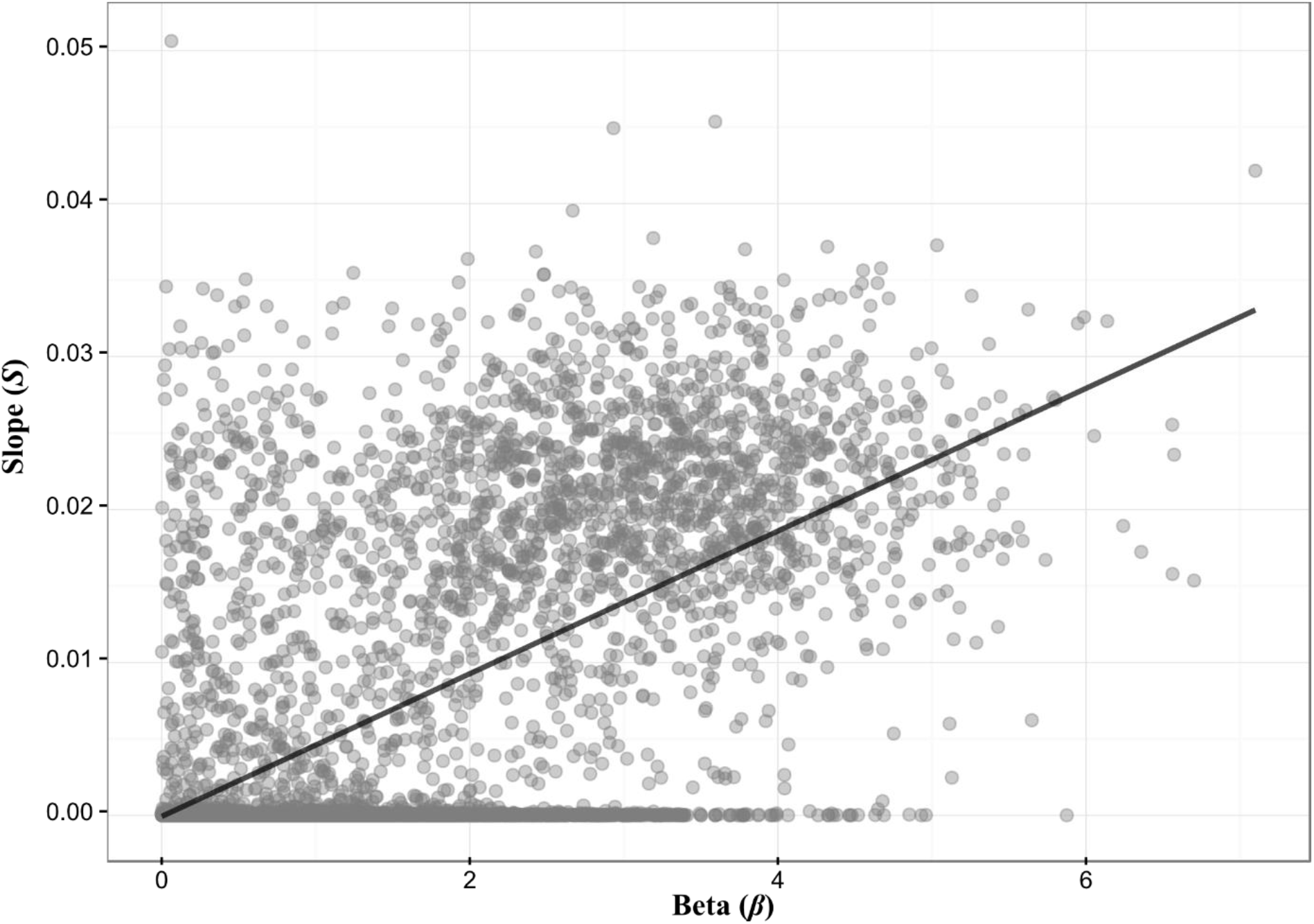
Correlation between the geographic cline slope parameter (*S*) and positive values of the genomic cline parameter *β*.

Moreover, we tried to connect genomic and geographic cline parameters with our estimated probability that individual loci belong to the small fraction of the genome showing the highest introgression rate. As expected, private SNPs located on the outer frame of the JAFS, for which we inferred a small probability of introgression, were usually associated with steep cline slopes. By contrast, loci occupying the most central part of the JAFS, which were assigned the highest introgression probabilities, were also characterized by shallower geographic clines. Finally, when restricting the analysis to the 5% of loci with the highest introgression probability, we show that a majority of their geographic cline centres are located outside of the contact zone, with a spatial shift more pronounced in the *S. aegyptiaca* direction (Fig. 7a). Similarly, the distribution of the genomic cline parameter *α* also showed a deficit of values around zero, and a majority of loci with an excess of *S. senegalensis* ancestry, in keeping with the general asymmetry of the exchanges between the two species (Fig. 7b).

**Figure 7:**
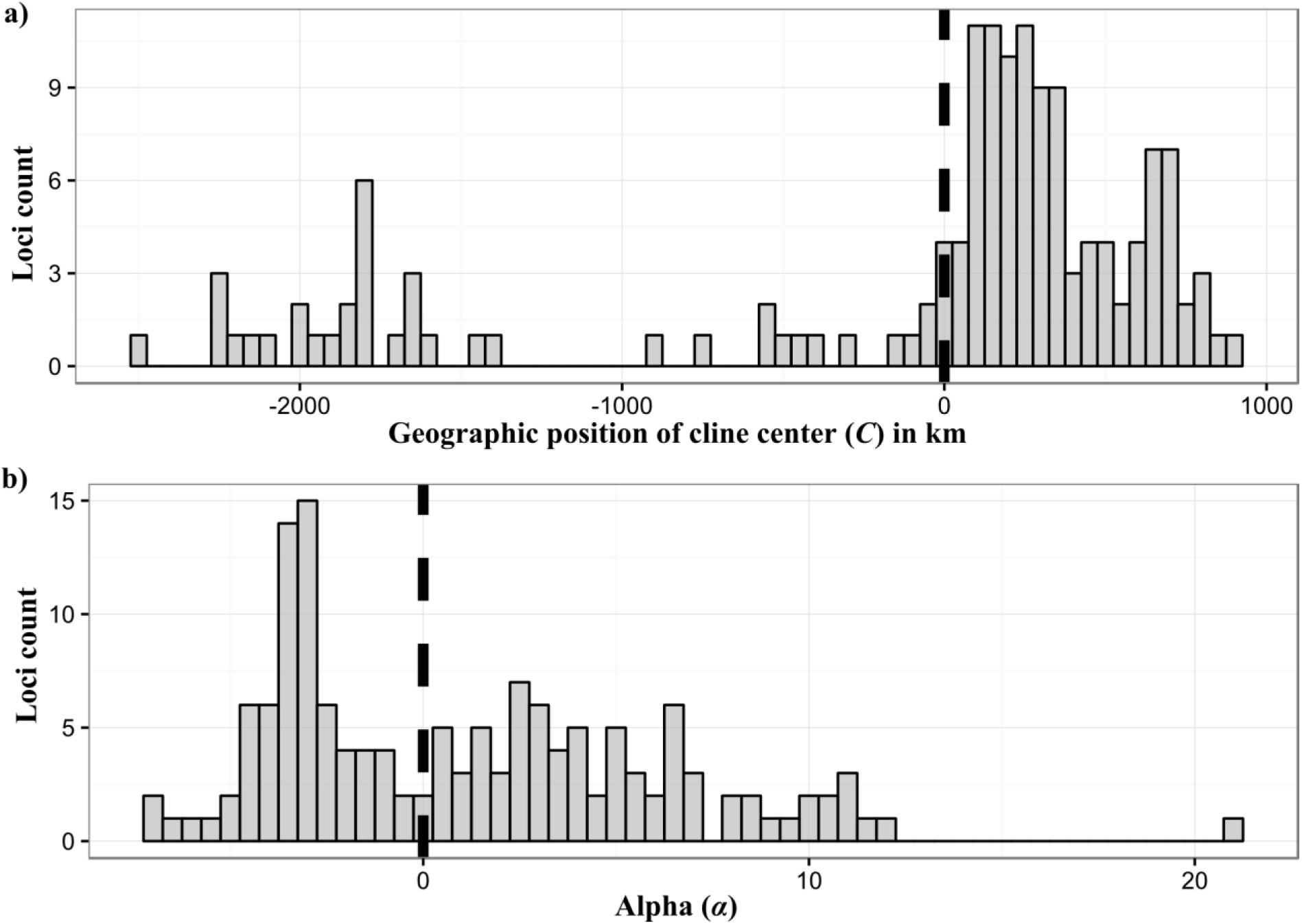
Geographic distribution of **a)** the geographic cline center (*C*) and **b)** the genomic cline parameter *α* for loci showing a probability of introgression greater or equal to 95%.

## DISCUSSION

### Divergence history and semi-permeability to gene flow

Our results using high-throughput genotyping confirmed previous observations, based on a limited number of nuclear markers, that the flatfish *S. senegalensis* and *S. aegyptiaca* are genetically divergent sibling species, which are still exchanging genes across their natural hybrid zone near Bizerte lagoon (She *et al*., 1987; Ouanes *et al*., 2011; Souissi *et al*., 2017). However, the proportion of hybrid genotypes detected here in the contact zone (i.e. 3 among 64 individuals) is much lower than established in previous studies. Although this could be due to large temporal variations in the rate of hybridisation, the most plausible hypothesis is that real hybrids (e.g. F1, F2 and first generations backcrosses) are more easily distinguished from introgressed genotypes using a high number of diagnostic markers (e.g. Pujolar *et al*., 2014; Jeffery *et al*., 2017). Our results thus support the existence of an abrupt geographic transition between the two species across their contact zone, where predominantly parental genotypes occur in sympatry with an overall low frequency of hybrids. These observations are consistent with the tension zone model (Barton and Hewitt, 1985), in which the hybrid zone is maintained by a balance between the influx of parental genotypes from outside the zone and the counter-selection of hybrid genotypes inside the zone. This interpretation is also in keeping with the finding of transgressive body shape variation attributed to a reduced condition index of admixed genotypes from the contact zone (Souissi *et al*., 2017).

The regions located outside the contact zone, which contain only parental genotypes, provided relevant information to infer the demo-genetic history of divergence between these two species. Gene exchange between *S. senegalensis* and *S. aegyptiaca* is best explained by a secondary contact model with heterogeneous rates of introgression along the genome. Similar divergence histories have already been found between geographical lineages or ecotypes of the same species, like in the European sea bass and anchovy (Tine *et al*., 2014; Le Moan *et al*., 2016). Using the same approach as implemented here, these studies could determine that a minor but significant fraction of the genome (i.e. 20 to 35%) does not neutrally introgress between hybridizing lineages/ecotypes, thus providing a quantitative assessment of the degree of semi-permeability to gene flow. In the present case, by contrast, we found that the great majority of the genome (approximately 95%) experiences a highly reduced effective migration rate (i.e. < 1 migrant per generation) between species. Moreover, the inferred divergence period estimated using a mutation rate of 10^-8^ per bp per year and a generation time of 3 to 5 years corresponds to a separation time of *ca*. 1.1 to 1.8 Myrs, which together with the mitochondrial sequence divergence of 2%, indicates a relatively ancient speciation event. This timing of divergence is much older than the approximately 300 Kyrs that were inferred between glacial lineages in anchovy and sea bass (Tine *et al*., 2014; Le Moan *et al*., 2016). Therefore, our results are consistent with the prediction that more anciently diverged species should display stronger reproductive isolation due to the establishment of more genetic barriers (Orr, 1995; Moyle and Nakazato, 2010; Roux *et al*., 2016), which is reflected here by a lower level of permeability to gene flow compared to sea bass and anchovy.

Another aspect of this low miscibility of the two genomes was captured by individual locus cline parameters. Consistent with the view that stepped clines generate a strong barrier to gene flow (Szymura and Barton, 1986), we found a high proportion of loci combining extremely low introgression rates (i.e. positive *β* outliers) and steep geographic clines (i.e. cline width below the average) with centres colocalized in the central part of the hybrid zone. Because this type of situation generates strong linkage disequilibrium, each locus is expected to receive, in addition to its own selective coefficient, the effect of selection on other loci (Barton, 1983; Kruuk *et al*., 1999). This makes the genome acting almost as a single underdominant locus causing a global reduction in gene flow, which sometimes is referred to as a “congealed genome” (Turner, 1967; Bierne *et al*., 2011; Gompert *et al*., 2012b). In contrast to this genome-wide reduction in gene flow, we also found support for a small proportion of genomic regions with higher-than-average introgression rates between species.

### Multiple approaches to detect increased introgression rates

Differential introgression among loci was evidenced by combining several approaches that capture different but complementary aspects of the data. For instance, genomic and geographic cline methods both rely on the sampling of admixed genotypes from the hybrid zone, whereas the modelling approach based on the JAFS relies of the joint distribution of allele frequencies between parental populations located outside of the hybrid zone. By combining these methods, we found that the 5% of loci with higher-than-average introgression rates identified with the JASF approach generally display shifts in both geographic and genomic cline centres. Therefore, our method to detect increased introgression while accounting for the demographic divergence history identifies genomic regions that also depart from the genome average in those places where the two species meet and admix.

Discordant clines, were not symmetrically distributed between the *S. senegalensis* and *S. aegyptiaca* side of the hybrid zone. The majority of the highly introgressed loci were shifted into the *S. aegyptiaca* geographic range. For these loci, the excess of *S. senegalensis* ancestry (measured by the genomic cline parameter *α*) was positively related to the extent of the spatial shift of the cline center into the *S. aegyptiaca* territory. This correlation was much stronger than previously observed in a bird hybrid zone study (Grossen *et al*., 2016), possibly due to a stronger variance in cline shift in the *Solea* system. By contrast, much fewer loci were found with increased introgression rates in the opposite direction. Such loci generally displayed cline centres shifted far away from the contact zone, near the entrance of the Mediterranean Sea. The spatial shift in cline center was in this case only weakly positively correlated with the excess of *S. aegyptiaca* ancestry. This could be expected, however, since the genomic cline method has limited power to infer cline parameters outside the range of observed hybrid indexes in admixed genotypes (Gompert *et al*., 2012a), which in our case were biased in favour of *S. aegyptiaca* ancestry.

### Differential introgression patterns and the dynamics of the hybrid zone

The inferred history of divergence between *S. senegalensis* and *S. aegyptiaca* places the beginning of gene flow *ca*. 18 to 30 Kyrs ago, supporting a scenario of secondary contact at the end of the last glacial period. Therefore, the hybrid zone is probably sufficiently old for the dynamics of introgression to be well established. This supports the idea that reproductive isolation between *S. senegalensis* and *S. aegyptiaca* was strong enough to prevent genetic homogenization throughout most of the genome during this period. At the same time, the small fraction of introgressing loci that were still able to cross the species barrier indicate the existence of genomic regions unlinked to reproductive isolation loci, either due to a local lack of barrier loci and/or to a high local recombination rate (Roux *et al*., 2013).

Whether pre- or postzygotic effects could explain the observed asymmetric introgression under the hypothesis of a stable hybrid zone remains to be fully addressed. Alternatively, the geographic pattern of asymmetrical introgression evidenced here could reveal a movement of the hybrid zone (Buggs, 2007), especially if the markers concerned are randomly spread across the genome (Wielstra *et al*., 2017). In the absence of a well-assembled reference genome, the genomic distribution of introgressing loci could not be addressed. However, a northward movement of the hybrid zone has already been proposed based on a temporal trend of increasing abundance of *S. aegyptiaca* in Bizerte lagoon (Ouanes *et al*., 2011). Therefore, our results probably reflect a recent or ongoing shift of the hybrid zone, with *S. aegyptiaca* incorporating the small fraction of compatible genes from *S. senegalensis* as it moves northward in its territory, leaving behind a tail of neutral introgressed alleles. Similar signatures have already been reported in other sister species such as in house mice (Wang *et al*., 2011), rabbits (Carneiro *et al*., 2013), salamanders (Visser *et al*., 2017), toads (Arntzen *et al*., 2017), newts (Wielstra *et al*., 2017) and chikadees (Taylor *et al*., 2014). The possible causes of hybrid zone movement include the tracking of environmental changes, fitness differences among individuals from the two species, or gradients of population density (Barton and Hewitt, 1985; Buggs, 2007; Taylor *et al*., 2015; Gompert *et al*., 2017). Future studies will have to establish which of these hypotheses accounts for the asymmetric introgression pattern in soles.

Opposite to the preferential direction of introgression, we also found a few loci within the *S. senegalensis* genetic background whose geographic center was apparently shifted near the entrance of the Mediterranean, far away from the contact zone. This pattern was also captured by the first and third PCA axes, along which we observed a slight differentiation between the *S. senegalensis* samples from Dakar, Cadiz and Annaba, which seems to be congruent with increased introgression in Mediterranean compared to Atlantic *S. senegalensis* samples. Long range gene flow is unlikely to explain these discordant clines that only concern a few loci. A more probable hypothesis would be that such loci exhibiting a shifted center could be related to an adaptive introgression of *S. aegyptiaca* alleles into the *S. senegalensis* background. Evidence for adaptive alleles spreading between species has been found in recent hybrid zones induced by human introductions (Fitzpatrick *et al*., 2010), but should be difficult to observe in historical hybrid zones where advantageous alleles have had ample time to spread (Hewitt, 1988). Strong barriers to gene flow only have a delaying effect on the dynamics of introgression of favourable alleles (Barton, 1979). If the barrier is strong enough, many generations may be required for advantageous alleles to extricate themselves from the hybrid zone, and therefore spatial allele frequency patterns related to adaptive introgression may appear even well after secondary contact. Moreover, the entrance of the Mediterranean Sea is a well-known barrier to dispersal for many marine organisms (Patarnello *et al*., 2007), which may delay the wave of advance of adaptive introgression clines as they travel towards Dakar to the south of *S. senegalensis*’ range. Alternatively, these clines may correspond to alleles of *S. aegyptiaca* ancestry, providing a local advantage to the *S. senegalensis* populations in the Mediterranean environment but not in the Atlantic. The hypothesis of adaptive introgression needs to be scrutinized in more details by focusing on the chromosomal signature of differentiation around the putative selected locus (Bierne, 2010).

## Conclusion

To conclude, our combination of analytical approaches provides new insights into the genomic architecture and the dynamics of gene flow between two highly divergent but still interacting parapatric species. Despite a relatively modest geographic coverage and the scarcity of available admixed genotypes in the tension zone, the genome-wide analyses taking into account the inferred history of divergence provided an efficient way to detect loci with deviant introgression behaviours. This is complementary to classical approaches based on genomic and geographic cline analyses. Our results bring new support for the tension zone model against a simple coexistence in sympatry. We show that differential gene flow has shaped genetic divergence across the tension zone, although most loci behave as if they were sitting on a congealed genome, rendering very unlikely the future remixing of the two genes pools via the creation of a hybrid swarm. Nevertheless, a few genes seem able to escape counter-selection, possibly due to different underlying processes. The first involves a possible movement of the zone, in which a shift of species range boundaries leaves a tail of neutral introgressed alleles behind the front of the invading species. The second is possibly related to the spread of globally or locally adaptive alleles into the range of the invaded species. Our results thus provide a snapshot on the genetic outcome of evolutionary processes potentially involved when divergent gene pools come back into contact after a long period of geographical isolation.

## Acknowledgments

This work was conducted during the PhD of A. Souissi with the financial support of programme PHC Maghreb No. XE27961 from the French Ministère des Affaires Etrangères to F. Bonhomme and Ministère de 1’Enseignement Supérieur et de la Recherche to L. B. Sfar. The authors are indebted to M.-T. Augé for skillful technical assistance.

## Data Archiving

Raw sequences data and VCF files will be deposed upon acceptance on Genbank and Dryad

